# A Nanoemulsion as an Effective Treatment Against Human Pathogenic Fungi

**DOI:** 10.1101/737767

**Authors:** Alexis Garcia, Yong Yi Fan, Sandeep Vellanki, Eun Young Huh, DiFernando Vanegas, Su He Wang, Soo Chan Lee

**Affiliations:** South Texas Center for Emerging Infectious Diseases (STCEID), Department of Biology, the University of Texas at San Antonio, San Antonio, TX 78249, USA; Michigan Nanotechnology Institute for Medicine & Biological Sciences, University of Michigan, Ann Arbor, MI 48109, USA; Department of Internal Medicine, University of Michigan Medical Center, Ann Arbor, MI 48109, USA

## Abstract

The emergence of immunocompromising diseases such as HIV/AIDS or other immunosuppressive medical conditions have opened an opportunity for fungal infections to afflict patients globally. An increase antifungal drug resistant fungi have posed a serious threat to patients. Combining these circumstances with a limited variety of antifungal drugs available to treat patients has left us in a situation where we need to develop new therapeutic approaches that are less prone to development of resistance by pathogenic fungi. In this study we present the utilization of the nanoemulsion NB-201 to control human pathogenic fungi. We found that the NB-201 exhibited in vitro activity against *C. albicans*, including both planktonic growth and biofilms. Furthermore, treatments with NB-201 significantly reduced the fungal burden at the infection site and presented enhanced healing process after subcutaneous infections by multidrug resistant *C. albicans* in a murine host system. NB-201 also exhibited in vitro growth inhibition activity against other fungal pathogens, including *Cryptococcus* spp, *Aspergillus fumigatus*, and Mucorales. Due to the nature of the activity of this nanoemulsion, there is a minimized chance of drug resistance to develop, thus presents a novel treatment to control fungal wound or skin infections.

## Introduction

During the past decade there has been an exponential growth in discoveries and medical advances for the treatment of human disease. This has led to better treatment for patients, and as a result we have been able to prolong human life. While these recent medical advances have certainly been beneficial overall, procedures such as solid organ transplants and cancer treatments have left many patients in an immunocompromised state. The emergence of immunocompromising diseases such as HIV/AIDS or other immunosuppressive medical conditions have opened an opportunity for fungal infections to plague patients globally (1–4).

*Candida albicans* is a human commensal fungus found on the skin, mucosal membranes, and the normal gut flora (5, 6). *C. albicans* is known to be an opportunistic fungus and the most common fungal pathogen which typically infects immunocompromised patients (3, 4). Treatment for candidiasis currently relies on three major classes of antifungal drugs including echinocandins, azoles, and polyenes (7, 8). Prior to the introduction of echinocandins, fluconazole was the most common drug used to treat *C. albicans* infections (7). Recently there has been an increase in cases of drug resistant *C. albicans* infections resulting in an increase in morbidity and mortality of patients (4, 9–11). One explanation for drug resistance is the development of mutations in the target genes of the antifungal drug (9). Secondly the overexpression of efflux pumps and multi-drug resistance genes could also lead to antifungal drug resistance (9). In addition, pathogenic fungi can also form biofilms that are resistant to antifungal drugs (12, 13). Thus, it is of upmost importance to develop new therapeutic approaches that are less prone to the development of resistance by pathogenic fungi.

Membrane disruptive nanoemulsions have been developed to control pathogenic bacteria (14, 15). One example is the nanoemulsion NB-201, which is an emulsification of refined soybean oil, water, glycerol, EDTA, Tween 20, and the surfactant benzalkonium chloride (BZK), which is commonly used as an antimicrobial preservative in drugs, topical antiseptic, clinical disinfectant, and as a sanitation agent in the food industry (14, 16–18). The NB-201 formulation exhibited in vitro growth inhibition activity against *Pseudomonas aeruginosa* and in vivo activity in a burnt wound animal model. Treatment of animals infected with *Staphylococcus aureus* with NB-201 by using a murine skin abrasion wound model demonstrated the therapeutic potential of this nanoemulsion formulation (19, 20). NB-201 has also shown activity against Multi-Drug Resistant S. *aureus* (MRSA) in a study using a porcine burnt wound model (14). The NB-201 formulation is also able to greatly reduce the inflammation of infected wounds and is nontoxic to the skin making it a viable method of treatment for topical bacterial infections (14).

In this study we present the utilization of NB-201 to control pathogenic fungi. NB-201 exhibited in vitro efficacy against *C. albicans* planktonic cells and biofilms. In addition, NB-201 treatments significantly reduced fungal burden at the infection site and presented enhanced healing process after subcutaneous infections by *C. albicans* in a murine host system. NB-201 also exhibited in vitro growth inhibition activity against other fungal pathogens, including *Cryptococcus* spp, *Aspergillus fumigatus*, and Mucorales. Fungal susceptibility against NB-201 was observed regardless of antifungal drug resistance. Due to the nature of the activity the nanoemulsion presents, there is a minimized chance of drug resistance to develop, thus presenting a novel way to control fungal wound or skin infections

## Results

### In vitro activity against *Candida albicans* planktonic cells and pre-formed biofilms

*C. albicans* isolates including antifungal drug resistant strains (Table 1) were tested. 96-well plates were inoculated with each *C. albicans* strain and the NB-201 nanoemulsion (NE) (10%) added in ratios ranging from 1:1-1:2048. Minimum inhibitory concentrations (MICs) were determined by using a 100% killing point of the *C. albicans* planktonic cells collected at 1, 24, 48, and 72-hour post addition of NB-210 to the media (Table 2). We observed within 1 hour, a concentration of 1:512 of the NE was able to kill all the planktonic cells plated. As the incubation time was increased, we observed a lower MIC was required. At 24 hours, a concentration of 1:1024 was able to kill all the strains plated. Within 48 hours the concentration of NB-201 required to kill all ten strains remained at an MIC of 1:1024. At 72 hours the MIC for 100% killing of the strains plated was lowered to a concentration of 1:2048 (Table 2).

**Table 1.** fungal pathogens used in this study.

| Species | Strain | Info |
| --- | --- | --- |
| *Candida albicans* | SC5314 | Wild type |
| *Candida albicans* | TW1 | Clinical isolate (No Mutation) |
| *Candida albicans* | TW2 | Drug resistance observed (MDR1) |
| *Candida albicans* | TW3 | Drug resistance observed (MDR1) |
| *Candida albicans* | TW17 | Multi drug resistance observed (CDR1, MDR1, ERG11) |
| *Candida albicans* | 4639 | F449S, T229A (Erg11p substitutions) MDR1, CDR1 |
| *Candida albicans* | 6482 | D116E, K128T, Y132H, D278N, G464S, P230L (Erg11p substitutions) |
| *Candida albicans* | 4617 | F449S, T229A (Erg11p substitutions) |
| *Candida albicans* | 3731 | F126L, K143R (Erg11p substitutions) MDR1 |
| *Candida albicans* | 2240 | V437I (Erg11p substitutions) MDR1, ERG11 |
| *Candida albicans* | 412 | K128T (Erg11p substitution) |
| *Cryptococcus neoformans* | H99 | Serotype A |
| *Cryptococcus neoformans* | R265 | Serotype B |
| *Cryptococcus neoformans* | WSA87 | Serotype C |
| *Cryptococcus neoformans* | R4247 | Serotype D |
| *Aspergillus fumigatus* | CEA10 | Wild type, MAT1-1, clinical isolate |
| *Aspergillus fumigatus* | SRRC 2006 | Reference strain for identification and morphological observation (ICPA) |
| *Aspergillus fumigatus* | V044-58 | Azole resistant clinical isolate |
| *Aspergillus fumigatus* | F14946 | Azole resistant clinical isolate |
| *Aspergillus fumigatus* | F13747 | Azole resistant clinical isolate |
| *Aspergillus fumigatus* | F14532 | Azole resistant clinical isolate |
| *Aspergillus fumigatus* | F13746 | Azole resistant clinical isolate |
| *Aspergillus fumigatus* | F12776 | Azole resistant clinical isolate |
| *Aspergillus fumigatus* | F14403 | Azole resistant clinical isolate |
| *Aspergillus fumigatus* | F6919 | Azole resistant clinical isolate |
| *Aspergillus fumigatus* | F16216 | Azole resistant clinical isolate |
| *Rhizopus delemar* | DI18-58 | Drug resistant clinical isolate |
| *Rhizopus delemar* | DI18-59 | Drug resistant clinical isolate |
| *Rhizopus microsporus* | DI18-62 | Drug resistant clinical isolate |
| *Rhizopus microsporus* | DI18-63 | Drug resistant clinical isolate |
| *Mucor circinelloides* | DI18-65 | Drug resistant clinical isolate |
| *Mucor circinelloides* | DI18-66 | Drug resistant clinical isolate |
| *Cunninghamella* | DI18-68 | Drug resistant clinical isolate |
| *Cunninghamella* | DI18-69 | Drug resistant clinical isolate |
| *Lithemia* | DI18-71 | Drug resistant clinical isolate |
| *Lithemia* | DI18-72 | Drug resistant clinical isolate |

**Table 2.**
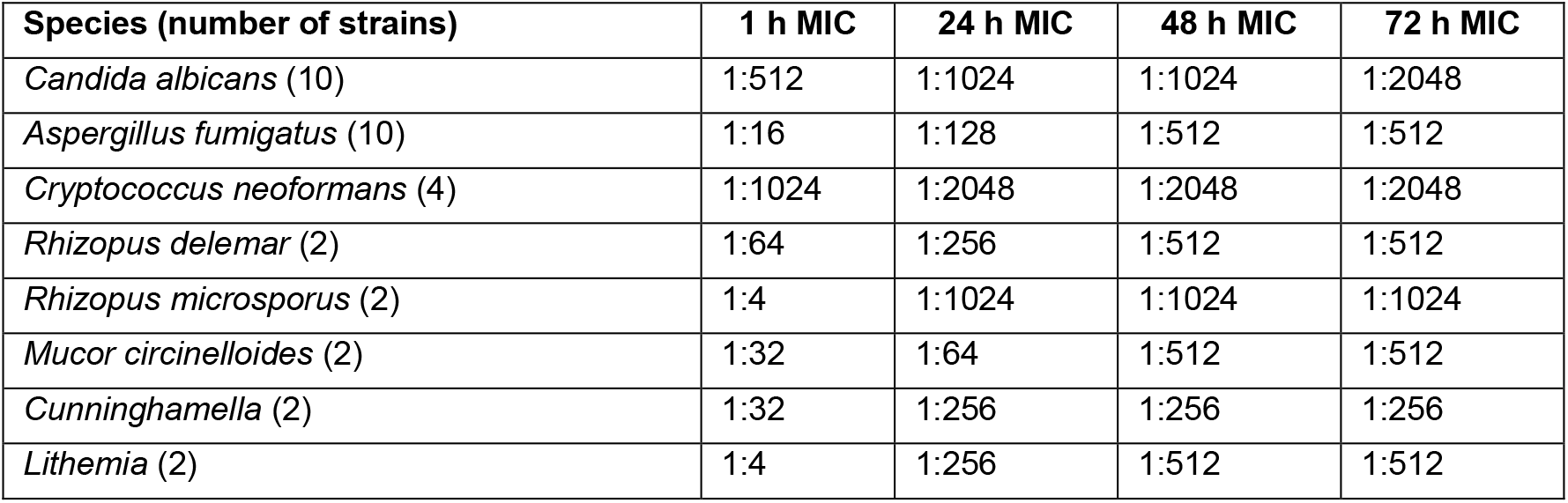
*In vitro* activity of NB-201

The ability of *C. albicans* to form biofilms, which increases antifungal drug resistance, is a major virulence factor observed in the clinical setting (12, 13). To test the efficacy of NB-201 on *C. albicans* biofilms, two multi-drug resistant clinical isolates TW1 and TW17 (21) were chosen. The *C. albicans* clinical isolates TW1 and TW17 were plated on 96-well plates and allowed to form a biofilm over the course of 24 hours (PFB). We then treated these PFBs with the NE added in various ratios ranging from 1:1-1:2048 followed by a second-generation tetrazolium (XTT) metabolic assay (Sigma-Aldrich) (22) to measure ratio of the metabolism, indicative of disruptions of the biofilms, after NB-201 treatments. Within 2 hours a NE concentration of 1:32 was able to inhibit 100% of the metabolism in the TW1 clinical isolate (Figure 1 A). At 4 hours, a concentration of 1:64 was inhibiting greater than 50% of the metabolism in the PFBs (Figure 1 B), while incubation with NB-201 at 6 hours exhibited higher inhibition up to 90% at the same concentration (Figure 1 C). At 24 hours, a concentration of 1:128 presented 90% inhibition of metabolism in the TW1 PFBs (Figure 1 D) while a concentration of 1:256 presented 75% metabolism inhibition at 48 hours (Figure 1 E). After 72 hours of exposure to NB-201, a concentration of 1:256 presented 85% inhibition of the metabolism of the PFBs (Figure 1 F). A similar trend was observed with the TW17 isolate. Within 2 hours post addition of NB-201, a concentration of 1:16 was required for 100% metabolism inhibition, while we find it important to note that a concentration of 1:32 inhibited 95% of the metabolism in the PFBs (Figure 1 G). Within 4 hours of exposure to the NE the PFBs presented 100% metabolism inhibition at a 1:32 concentration with >50% inhibition being observed at a concentration of 1:64 (Figure 1H). After 6 hours of exposure, a 1:64 concentration of NB-201 inhibited 80% of the metabolism in the PFBs (Figure 1I). A 24-hour exposure to the NE at a concentration of 1:128 presented a 70% inhibition of the metabolism in the TW17 PFBs while a concentration of 1:64 inhibited 100% of the metabolism (Figure 1J). At 48 hours of exposure a concentration of 1:256 presented 85% inhibition of metabolism (Figure 1K) while a 72-hour exposure increased that to 100% inhibition at the same concentration (Figure 1L). A similar result was obtained to measure the efficacy of NB-201 in disruption of *C. ablcians* biofilms by using a crystal violet staining in the TW1 isolates (Supplementary Figure 1) and the TW17 isolates (Supplementary Figure 2). Two independent experiments were performed with a similar result with each experiment containing biological repeats.

**Figure 1.**
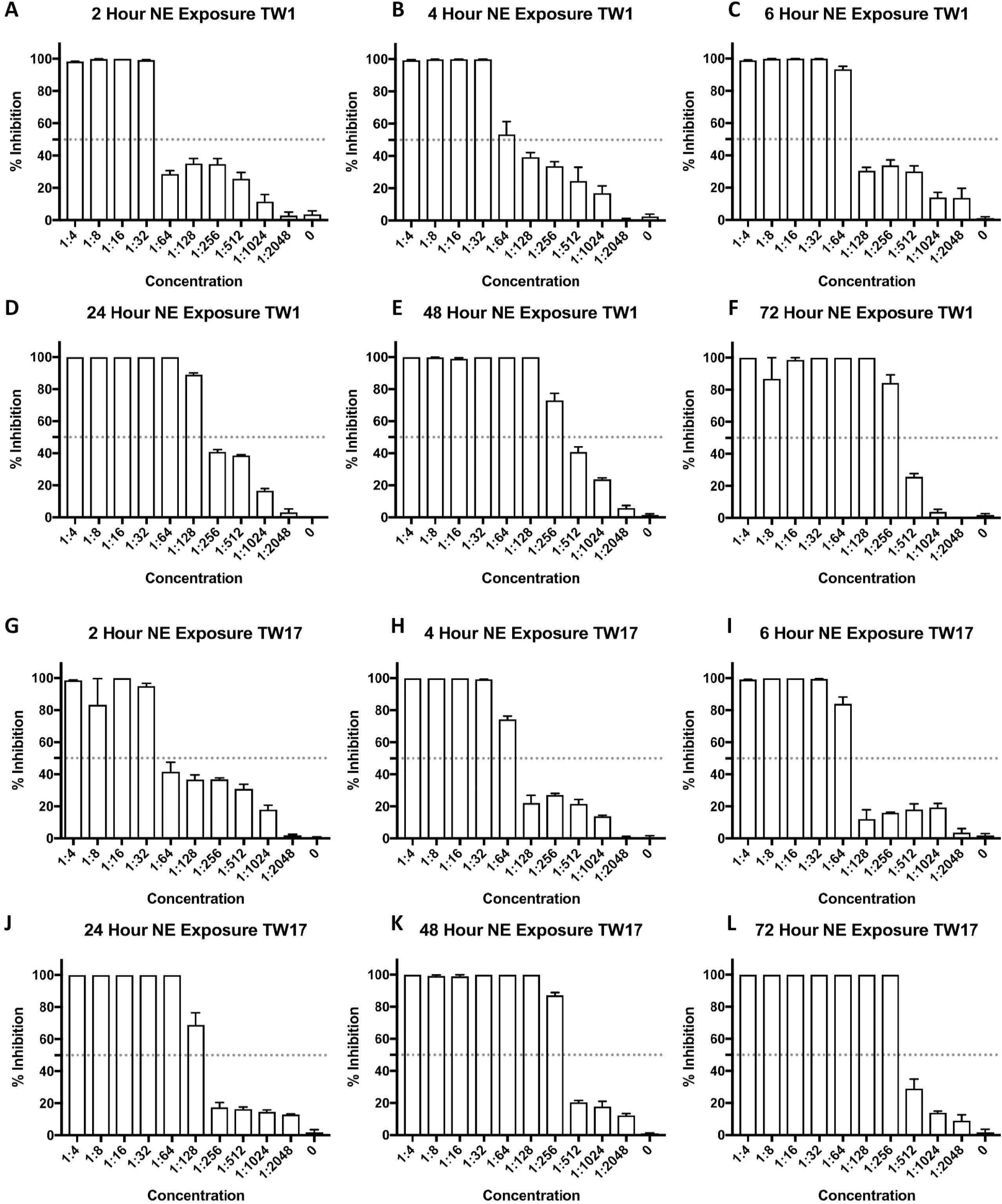
Measurement of Metabolism in *C. albicans* Drug Resistant TW1 Clinical Isolate Pre-Formed Biofilms. Multi-drug resistant *C. albicans* clinical isolates TW1 and TW17 were plated in a 96-well plate containing RPMI media at a concentration of 1×10^6^ and incubated for 24 hours to form a biofilm on the bottom of the wells. The media containing the nanoemulsion treatment was then removed followed by the biofilms being treated with XTT solution and quantified. (A) 2 hours post treatment with NB-201 (B) 4 hours post treatment with NB-201 (C) 6 hours post treatment with NB-201 (D) 24 hours post treatment with NB-201 (E) 48 hours post treatment with NB-201 (F) 72 hours post treatment with NB-201. (G) 2 hours post treatment with NB-201 (H) 4 hours post treatment with NB-201 (I) 6 hours post treatment with NB-201 (J) 24 hours post treatment with NB-201 (K) 48 hours post treatment with NB-201 (L) 72 hours post treatment with NB-201.

### In vivo activity of NB-201 against *Candida albicans* subcutaneous infection

In vivo efficacy of NB-201 was tested by using a murine subcutaneous infection model. Mice were infected subcutaneously with the multi-drug resistant *C. albicans* isolates TW1 or TW17. We then treated the mice with NB-201 via subcutaneous injection (see Material and Methods). After two days of treatment with the NB-201, we euthanized the mice and collected tissues at the site of infection. To measure fungal burden in the tissues, we plated the homogenized tissue onto YPD agar plates and counted the colony forming units (CFU). Treatment with NB-201 resulted in a significant decrease of fungal CFUs in mice infected with TW1 (P=0.013) and with TW17 (P=0.002) compared to PBS-treated control groups (Figure 2A). The *C. albicans* strains from the first infection experiment were recovered and used for a second round of the assay. In the group of mice infected with the recovered TW1 or TW17, we observed a significant decrease of CFUs in the tissues of mice treated with the NE (P=0.005 or P=0.004, respectively) when compared to mice treated with PBS. (Figure 2B). These results demonstrate that NB-201 has the efficacy to control *C. albicans* infections while the resistance and/or tolerance against NB-201 is less likely to develop. The azole fluconazole still remains as one of the first drugs administered for the treatment of candidiasis (23). With this in mind we wanted to compare the efficacy of NB-201 to that of fluconazole. In similar fashion to our previous experiment, we began by using a murine subcutaneous infection model. Mice were infected with either the wild-type *C. albicans* SC5314, or the fluconazole resistant strain TW17 (Figure 3) followed with treatment by either a mock injection (PBS), NB-201, or fluconazole over the course of 72 hours. Treatment of NB-201 shows a significant reduction in swelling and inflammation at the infection site when compared to treatment with fluconazole despite the WT strain being susceptible to fluconazole (Figure 3A). A similar result with the NB-201 treatment can be observed in the mice infected with the azole-resistant strain TW17 with the exception of fluconazole showing no signs of treatment (Figure 3B).

**Figure 2.**
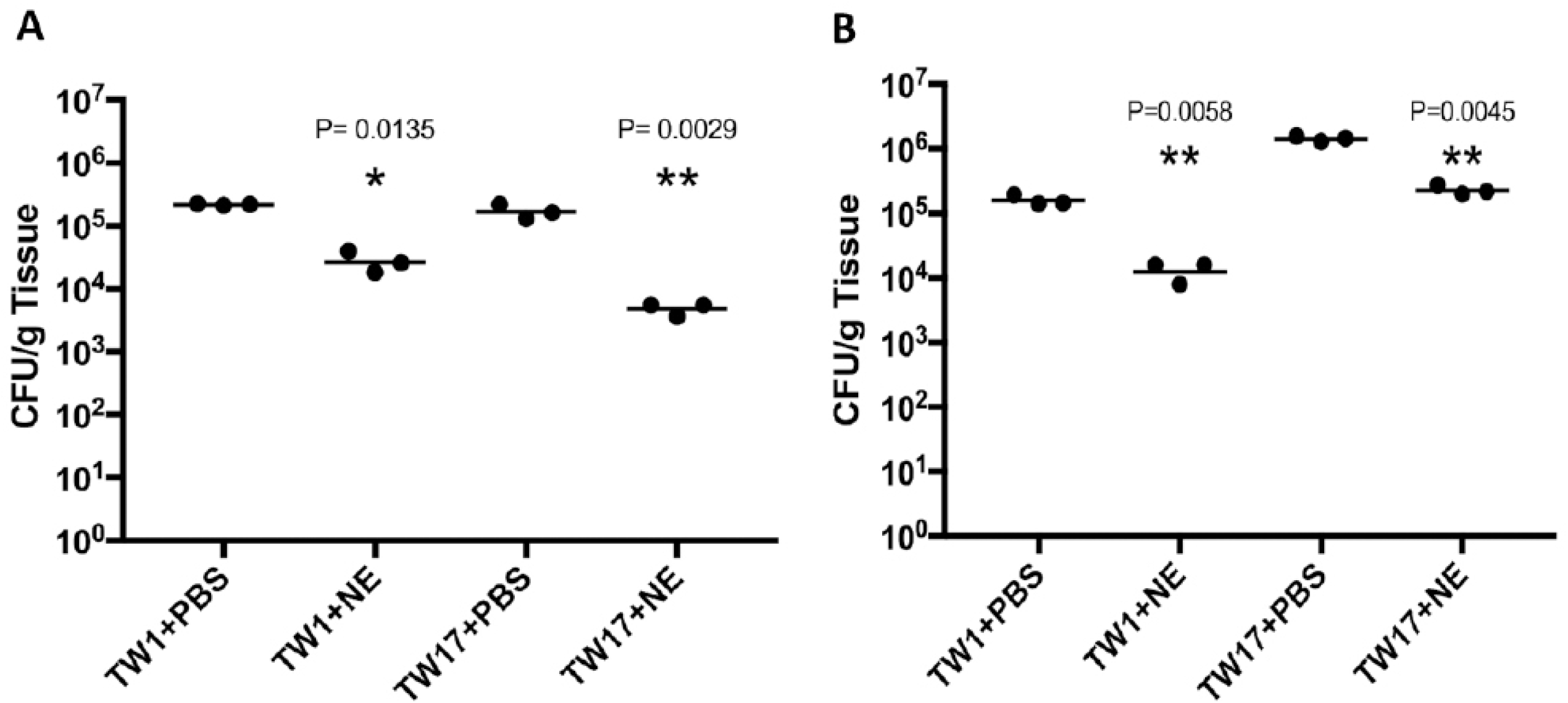
In Vivo Efficacy of NB-201 Via Subcutaneous Infection. In vivo efficacy of NB-201 was tested by using a murine subcutaneous infection model. Mice were infected subcutaneously with the multi-drug resistant *C. albicans* isolates TW1 or TW17 We then treated the mice with NB-201 via subcutaneous injection. To measure fungal burden in the tissues, we plated the homogenized tissue onto YPD agar plates and counted the colony forming units (CFU). (A) Initial infection. A significant reduction in fungal burden can be observed in mice treated with NE in both TW1 (P=0.0135) and TW17 (P=0.0029). (B) Recovered strain infection. Mice were infected with strains recovered from the initial infection to check for development of resistance to the NE. A significant reduction in the fungal burden can be observed in mice treated with NB-201 in both TW1 (P=0.0058) and TW17 (P=0.0045).

**Figure 3.**
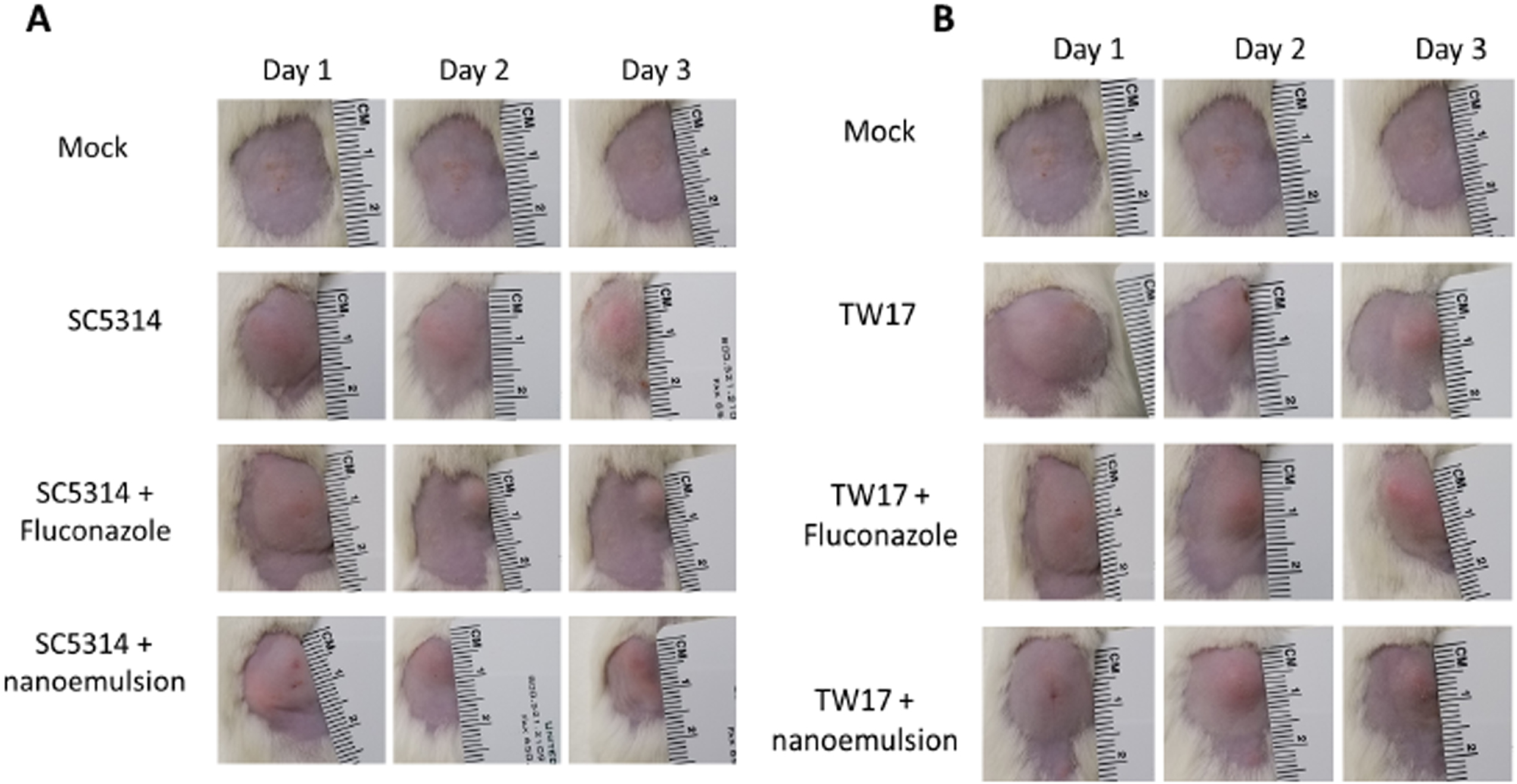
Comparison of NB-201 Efficacy to Fluconazole Via Subcutaneous Infection with wildtype *C. albicans.* Comparison of the in vivo efficacy of NB-201 compared to fluconazole was tested by using a murine subcutaneous infection model. Mice were infected subcutaneously with the wildtype *C. albicans* (SC5314), or with the fluconazole resistant *C. albicans* (TW17). Subsequent treatments via injection of either fluconazole or NB-201 we followed over 72 hrs. Due to SC5314 being susceptible to fluconazole, a reduction was observed in mice treated with the azole. When compared to the azole treated mice, NB-201 presented a greater healing ability by showing a reduction of swelling and inflammation over the course of 72 hrs. TW17 is intrinsically resistant to fluconazole, thus no reduction was observed in mice treated with the azole. When compared to the azole treated mice, NB-201 presented a greater reduction of swelling and inflammation over the course of 72 hrs.

Following our subcutaneous model, we took tissue samples from uninfected mice, mice infected with *C. albicans* TW17, and mice infected and treated with NB-201. We then stained our tissue samples with a haemotoxylin and eosin stain. When compared to the non-infected mice (Figure 4A), post *C. albicans* infection tissue presents a large collection of infiltration around the hair follicles, deep dermis, and superficial fat (Figure 4B). After treatment with NB-201 we can observe a reduction in the infiltration of cells within the deep dermis, superficial fat and hair follicles (Figure 4C). In the untreated tissue the layers of the skin are not as clearly defined when compared to the uninfected tissue. Treatment with NB-201 resulted in the layers of the skin to be more defined when compared to the untreated tissues. These observations further support the efficacy of the nanoemulsion to control multi-drug resistant *C. albicans* infections.

**Figure 4.**
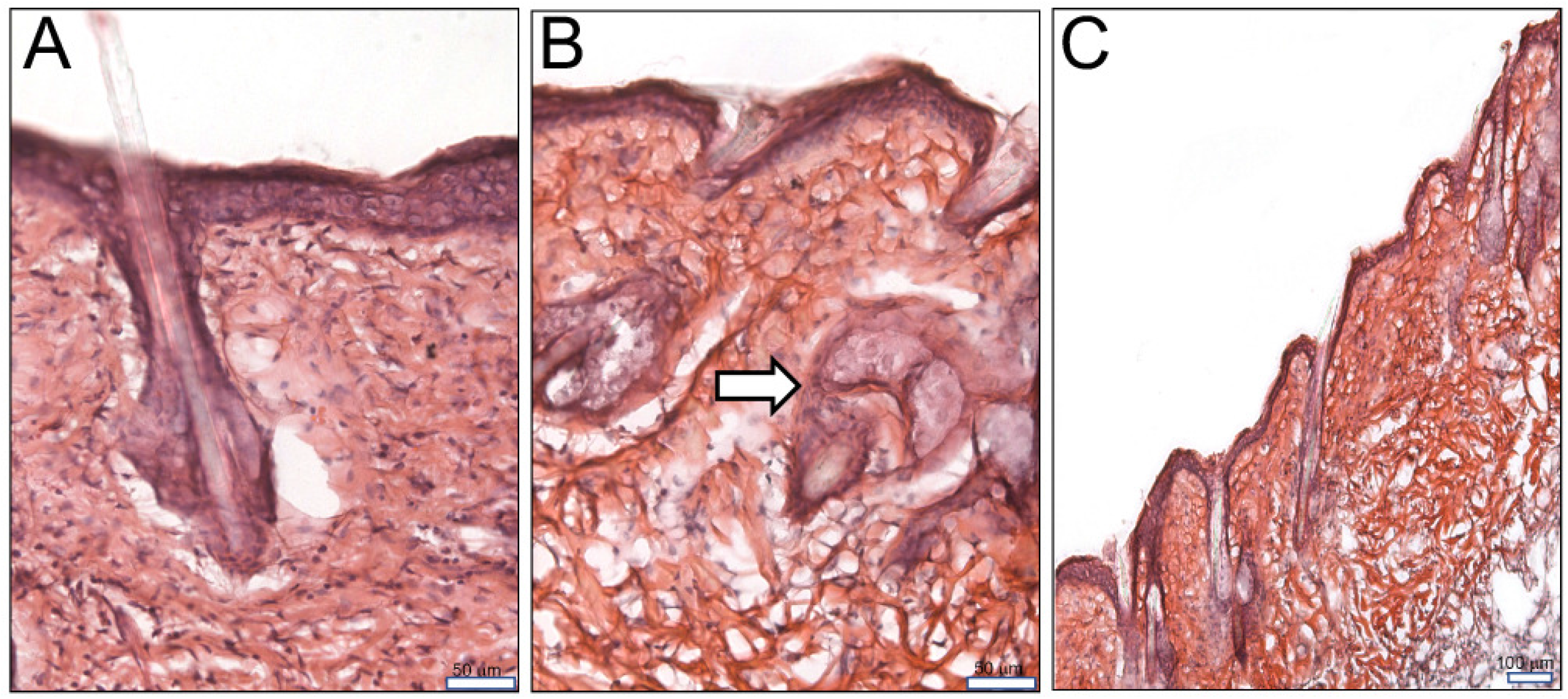
Histopathological Analysis of Mouse Skin Tissue Post-Subcutaneous Infection and Treatment with NB-201. Samples from uninfected mice, mice infected with *C. albicans*, and mice infected and treated with NB-201 were sectioned. We then stained our tissue samples with a haemotoxylin and eosin stain. (A) Uninfected mouse skin tissue. (B) Infected and untreated mouse skin tissue. Accumulation of infiltrates at the hair follicles can be observed (white arrow). (C) After treatment with NB-201 we can observe a reduction in the infiltration of cells within the deep dermis, superficial fat and hair follicles. Scales are 50 μm (A and B) or 100 μm.

### In vitro activity of NB-201 against other pathogenic fungi

The formulation of the NB-201 was further tested to examine the ability to kill other pathogenic fungi (Table 1) in a similar fashion to what was observed in *C. albicans.* We approached this by inoculating 96-well plates with varying fungal strains and added NB-201 in ratios ranging from 1:1-1:2048. We then measured the minimum inhibitory concentrations (MIC) by using a 100% killing point of the fungal strains collected at 1, 24, 48, and 72-hour post addition of NB-210 to the media (Table 2).

### Aspergillus fumigatus

We performed a checkerboard assay with ten different strains of *Aspergillus fumigatus* including drug resistant strains, all of which are known clinical isolates (Table 1). Within one hour we observed a concentration of 1:16 showed complete killing of all clinical isolates. We would like to note that a concentration of 1:128 was able to kill seven out of the ten *A. fumigatus* clinical isolates within the same timepoint (Table 2). As incubation time with the NE progressed, we observed a reduction in the MIC required to kill all of the *A. fumigatus* clinical isolates. Within 24 hours a concentration of 1:128 showed 100% killing of these clinical isolates (Table 2). Finally, at 48 and 72 hours all ten of the *A. fumigatus* clinical isolates were killed at a concentration of 1:512 (Table 2).

### Mucorales

We tested ten clinical isolates of varying *Mucorales* species (Table 1). One hour after incubation with NB-201 we observed a total MIC of 1:64 in *Rhizopus delemar* isolates, 1:4 in *R. microsporus* isolates, 1:32 in *Mucor circinelloides* isolates, 1:32 in *Cunninghamella* isolates, and 1:4 in *Lithemia* isolates (Table 2). In a similar fashion to what we have observed thus far, longer NE incubation with the fungi resulted in a lowered MIC. At 24 hours *R. delemar, Cunninghamella*, and *Lithemia* isolates presented an MIC of 1:256. *R. microsporus* isolates had an MIC of 1:1024, while *M. circinelloides* showed a MIC of 1:64 (Table 2). At 48 hours *R. delemar, M. circinelloides*, and *Lithemia* isolates resulted in a lowered MIC of 1:512. *Cunninghamella* isolates remained at 1:256 followed by *R. microsporus* presenting an unchanged MIC of 1:1024. At 72 hours we observe no changes in the MIC with any of the Mucorales strains (Table 2).

### Cryptococcus neoformans

Four different serotypes of *C. neoformans* against NB-201 were tested (Table 1). Within one hour an MIC of 1:1024 was able to kill all four serotypes of *C. neoformans* (Table 2). This was followed by 24, 48, and 72 hours showing an MIC of 1:2048 (Table 2). NE treated and untreated *C. neoformans* were plated in media containing propidium iodide, which is only able to penetrate the cellular membranes of dead or dying cells. Within 30 minutes of incubation a clear distinction between live and dead cells can be observed (Supplementary Figure 3 A) with the dead cells fluorescing red due to the propidium iodide stain (Supplementary Figure 3 B).

## Methods

### Fungal strains and growth conditions

The stains used in this study are listed in Table 1. *C. albicans* and *C. neoformans* strains were grown in liquid or solid yeast extract peptone dextrose [YPD, 10 g/L yeast extract, 20 g peptone, 20 g dextrose, 20 g agar (for plates only)] at 30°C. Mucorales were grown in potato dextrose agar (PDA, potato starch 4 g/L, dextrose 20 g/L, agar 15 g/L) or yeast extract peptone glucose agar (YPG, 3 g/L yeast extract, 10 g/L peptone, 20 g/L glucose, 2% agar, pH = 4.5) at 30°C in the light for four days. *A. fumigatus* strains were grown PDA at 30 °C for 4 days. To collect spores of Mucorales and *A. fumigatus*, sterile water (2 ml per plate) was added to the plate and spores were collected by gently scrapping the fungal mycelial mats.

### In vitro efficacy of NB-201 against *C. albicans* planktonic cells and biofilms

Initial concentration of NB-201 was 10% for all experiments in this study besides the data in supplementary table 1, where different concentration of NE was tested against *A. fumigatus.* The *C. albicans* strains were inoculated at a concentration of 1×10^6^ in a 96-well plate containing NB-201 serially diluted in RPMI (100 μl per well) ranged from 1:1 to 1:2048. 10μl samples taken at 1, 24, 48, and 72 hours from each well and plated on PDA agar plates, which were incubated for 48 hours. After incubation, every plate was examined for growth on the site of inoculation. For biofilms, *C. albicans* was plated at a concentration of 1×10^6^ in a 96-well plate containing RPMI media and incubated for 24 hours to form a biofilm on the bottom of the wells. After biofilm formation, the RPMI media was removed and the biofilms were washed with PBS. The media was then replaced with media containing NB-201 serially diluted in concentrations ranging from 1:1 to 1:2048. The media containing the nanoemulsion treatment was then removed at 2, 4, 6, 24, 48, and 72-hour time points. The biofilms were washed with PBS and stained with 0.6% crystal violet stain. The biofilms were then washed one more time with PBS to remove any residual crystal violet stain. Finally, the biofilms were de-stained with 33% acetic acid and 85 μl of the supernatant was transferred to a clean 96-well plate. The de-stained crystal violet supernatant was then read on a plate reader and quantified.

In similar fashion to the crystal violet assay *C. albicans* was plated in a 96-well plate containing RPMI media at a concentration of 1×10^6^ and incubated for 24 hours to form a biofilm on the bottom of the wells. After a biofilm was formed, the RPMI media was removed and the biofilms were washed with PBS. The media was then replaced with media containing NB-201 serially diluted in concentrations ranging from 1:1 to 1:2048. The media containing the nanoemulsion treatment was then removed at 2, 4, 6, 24, 48, and 72-hour time points and the biofilms were washed with PBS to remove any non-adherent cells. The biofilms were then treated with 100 μl of XTT solution containing 3.5 μl of menadione and incubated for 2 hours at 37°C. 85 μl of the supernatant was transferred to a clean 96-well plate and read in a plate reader then quantified. All in vitro efficacy experiments were repeated at least twice to verify the results.

### In vitro efficacy of NB-201 against Mucorales *Spp., C. neoformans*, and *A. fumigatus*

The respective fungal strains were inoculated at a concentration of 1×10^6^ in a 96-well plate containing NB-201 serially diluted in RPMI (100 μl per well). Dilution concentrations ranged from 1:1 to 1:2048. 10μl samples taken at 1, 24, 48, and 72 hours from each well and plated on PDA agar plates, which were incubated for 48 hours. After incubation, every plate was examined for growth on the site of inoculation.

### In vivo efficacy of NB-201 in a murine subcutaneous infection model

CD-4 mice weighing between 19-23 g were housed together. *C. albicans* strains SC5314, TW1, and TW17 were grown in YPD liquid media, washed in PBS, and suspended in PBS at a concentration of 1×10^6^. Under anesthesia, the dorsal fur of the mice was shaved. The exposed skin was washed with 70% ethanol and mice were infected with 1×10^6^ CFUs via subcutaneous injection on the shaved dorsal side. Subsequent subcutaneous injections of NB-201, PBS, or fluconazole followed at 6, 24, and 48 hours. The mice were euthanized at 72 hours, and the skin of the infected area was collected immediately for analysis.

The collected mouse tissue was placed in PBS on ice then homogenized with a tissue homogenizer. The homogenized tissue was then diluted 1:10 and plated on YPD agar plates treated with antibiotics to prevent unwanted bacterial growth. The plates were incubated at 32°C for 48 hours. The CFUs were then counted and quantified.

Animals were sacrificed and their skin tissue excised. The tissue samples were immersed in a formalin fixative agent. The tissue blocks were processed for cry-sectioning. 10-12 um micron thick sections were obtained with a cryostat and stained with Hematoxylin and Eosin for histopathological examination. Observations were made under light microscope, representative photomicrographs at 10X and 20X magnification were used for comparative study.

All murine experiments were conducted at the University of Texas at San Antonio (UTSA) in full compliance with all of the guidelines of the UTSA University Institutional Animal Care and Use Committee (IACUC) and in full compliance with the United States Animal Welfare Act (Public Law 98-198). The UTSA IACUC approved all of the murine studies under protocol number MU104-02-20. The experiments were conducted in the Division of Laboratory Animal Resources (DLAR) facilities that are accredited by the Association for Assessment and Accreditation of Laboratory Animal Care (AAALAC).

## Discussion

Previously, NB-201 presented a highly effective antimicrobial potential against various methicillin-resistant S. *aureus* strains both in vitro and in vivo (14). The surfactant in NB-201, benzalkonium chloride, is a known biocide found in many over-the-counter antibacterial handwipes, antiseptic creams, and other medically relevant consumer products (24). The use of this surfactant in NB-201 provides the benefit of minimizing harm to the human epidermis (25), as well as being an FDA approved biocide used in the clinical setting regularly. In our in vitro MIC experiments with *A. fumigatus* we found that the presence of BZK is required for NB-201 to function as a biocide. Furthermore, the presence of serum also presented an effect on the efficacy of NB-201s MIC (Supplementary Table 1). The in vitro susceptibility test with NB-201 was observed on every tested fungus, with an exceptional killing efficacy observed in all four serotypes of *C. neoformans.*

During our in vitro test we observed a similar trend in the efficacy of NB-201. Interestingly, we observed that longer incubation times with NB-201 resulted in a lowered MIC regardless of the fungal organism that was being tested. The top etymological agent of candidiasis, *C. albicans*, still ranks amongst the leading fungal organisms to cause infection in immunocompromised patients around the world and causes >50% of bloodstream infections in the US (3). The biofilms produced by this fungal organism makes it intrinsically harder to treat and is a growing problem and concern in the clinical setting (12, 13, 26, 27). We found that NB-201 has in vitro antifungal activity against planktonic form and biofilms of *C. albicans.* Furthermore, the in vitro activity was also observed against drug-resistant clinical isolates. In an animal subcutaneous infection model, NB-201 also exhibited antifungal activity against two azole-resistant strains, TW1 and TW17 (Figures 2, 3, and 4). These results demonstrate that NB-201 has anti-C. *albicans* activity both in in vitro and in vivo regardless of drug resistance. And C. albicans is less likely to develop resistant to NB-201 (Figure 2).

Our further in vitro data with other pathogenic fungi open possibility that NB-201 can be used for other fungal infections. When testing NB-201 against the four serotypes of *C. neoformans* we observed a dramatic killing ability. To further confirm what we observed, we employed a microscopy approach where we measured cell death with a propidium iodide stain. Within 30 minutes a clear distinction of live and dead cells was observed (supplementary figure 2). *C. neoformans* is the etiological component of cryptococcosis, an infectious fungal disease known to target the respiratory tract and central nervous system in humans (28–30). Exposure to *C. neoformans* is common amongst the general population, with majority of infectious cases resulting from reactivation due to latency in cell mediated immunity (30). Despite advances in modern medicine, the morbidity and mortality for *C. neoformans* infections remain unacceptably high with mortality rates of up to 20% in infected AIDS patients (28). *C. neoformans is resistant to the newest antifungal drug class echinodandin.* The infection mainly starts from the lungs and, therefore, the aerosol treatment of NB-201 could be applied especially to lung infections. Possible aerosol treatments with NB-201 could be explored in an in vivo model as a proof of concept for a novel form of treatment for respiratory cryptococcosis infection.

Known as one of the most prevalent airborne fungal pathogens in the world, *A. fumigatus* causes invasive aspergillosis (IA) in immunocompromised patients globally (31). In immunocompetent individuals IA is able to be naturally combated by the natural immunosuppressive abilities of the human body (31). One virulence factor presented by *A. fumigatus* is the ability to produce abounding numbers of spores with conidia resulting to be present in concentrations ranging from 1-100 conidia/m^−3^ in the air (31). Despite the high prevalence of IA in immunocompromised patients with a high morbidity and even higher mortality, not much is known about *A. fumigatus.* With this in mind we tested the efficacy of NB-201 in vitro with results similar to that observed in *C. albicans* (table 2) or methicillin-resistant S. *aureus* (14). Due to the large abundance of conidia present in the air *A. fumigatus* primarily infects the respiratory tract (32). Thus, in similar fashion to *C. neoformans* further exploring the efficacy of NB-201 in vivo would be of interest. This could be achieved via an intratracheal infection in a mouse model followed by subsequent treatments of NB-201 formulated as an aerosol.

Mucormycosis is a recently emerging opportunistic fungal infection (33). The typical causative agents for mucormycosis fall under the Mucorales family, which include *Rhizopus* spp., *Mucor* spp., and others (34). Due to the emergence of immunocompromising diseases such as HIV/AIDS or other medical conditions with lowered immunity there has been an increase in opportunity for fungal infections to afflict patients globally (35, 36). Mucormycosis presents itself with a mortality rate of ~50% in all mucormycosis cases and an unacceptable >90% mortality in disseminated mucormycosis cases (36–38). In the event that patients survive infection they typically suffer from disfiguration due to surgically removing infected tissue; a common way of treating mucormycosis infection aside from treatment with amphotericin B (39). Following the theme of which we have observed previously, NB-201 presented a killing ability comparable to that observed in our *C. albicans* in vitro experiments. Mucormycosis is typically caused when the acquired spores are inhaled into the body (36, 37). Although cutaneous infections of mucormycosis in patients undergoing traumatic injury have been reported (36, 40). It is in this type of cutaneous infection that NB-201 would present itself to be most effective in combating Mucorales infection.

Due to the nature of NB-201, the use of it as a topical treatment alternative for fungal infections with little to no drug resistance being developed by the fungi it is killing would be a possibility. Such treatments could include ointments for skin infections, aerosols for infection of the airways and lungs, and even oral washes for possible oral-pharyngeal infections.

## Conclusion

Currently, cases of immunocompromised patients being infected with antifungal drug resistant fungi have been rising. The development of drug resistance and the limited availability of antifungal drugs has left us in a scenario where we need to develop new therapeutic approaches that are less prone to the development of resistance by pathogenic fungi. Previously, NB-201 presented a highly effective antimicrobial potential against various methicillin-resistant *S. aureus* strains both in vitro and in vivo. In this study we have presented a novel use for the NB-201 nanoemultion formulation that presents killing abilities observed in vitro against 35 different fungi 30 of which are either clinical isolates or antifungal drug resistant strains. We also observed the reduction in inflammation, wound healing, and fungal pathogen clearing abilities of NB-201 in a murine host model (Figure 4). Due to the nature of the activity NB-201 presents, there is a minimized chance of drug resistance to develop, thus presenting a novel way to control fungal wound or skin infections.

## Acknowledgements

We are indebted to Jose Lopez-Ribot, David Denning, Bill Steinbach, Praveen Juvvadi, and Nathan Wiederhold for providing pathogenic fungal strains for this study. We are also indebted to the medical mycology group in UTSA for valuable discussions. We also thank Astrid Cardona for providing materials and tools for tissue histopathology. S.C.L. holds a Young Investigator Pilot Award in the Max and Minnie Tomerlin Voelcker Foundation. A.G. is supported by the UTSA RISE-PhD program (NIH/NIGMS RISE GM60655).

---

**Supplementary Table 1.** 1-hour *In vitro* activity of NB-201 at various BZK concentrations

| Nanoemulsion |  |  | MIC |  |
| --- | --- | --- | --- | --- |
| NE% | BZK% | EDTA% | No Serum | 25% Serum |
| 40 | 0.8 | 0.74 | 1:64 | 1:16 |
| 20 | 0.4 | 0.74 | 1:32 | 1:16 |
| 10 | 0.2 | 0.74 | 1:16 | 1:8 |
| 10 | 0 | 0 | No inhibition | No inhibition |

